# Plant genera *Cannabis* and *Humulus* share the same pair of well-differentiated sex chromosomes

**DOI:** 10.1101/2021.03.11.434957

**Authors:** D Prentout, N Stajner, A Cerenak, T Tricou, C Brochier-Armanet, J Jakse, J Käfer, GAB Marais

## Abstract

- We recently described, in *Cannabis sativa*, the oldest sex chromosome system documented so far in plants. Based on our estimate of its age, we predicted that it should be shared by its sister genus *Humulus*, which is known to also possess XY sex chromosomes.
- Here, we used transcriptome sequencing of a F1 family of *Humulus lupulus* to identify and study the sex chromosomes in this species using the probabilistic method SEX-DETector.
- We identified 265 sex-linked genes in *H. lupulus*, located on the chromosome that is also the *C. sativa* sex chromosome pair. Using phylogenies of sex-linked genes, we show that a region of these chromosomes had already stopped recombining in the common ancestor of the two species. Furthermore, as in *C. sativa*, Y gene expression was reduced in correlation to the position on the X chromosome, and strongly Y degenerated genes showed dosage compensation.
- Here we report, for the first time in the Angiosperms, a sex chromosome system that is shared by two different genera. Recombination suppression started at least 21-25 My ago, and then (either gradually or step-wise) spread to a large part of the sex chromosomes, leading to a strongly degenerated Y.

## Introduction

Among more than 15,000 dioecious angiosperm species (*i*.*e*. species with separate sexes; Renner, 2014), less than twenty sex chromosome systems have been studied with genomic data (Ming *et al*., 2011; Baránková *et al*., 2020). Most plants with sex chromosomes exhibit male heterogamety, with XY chromosomes in males, and XX chromosomes in females (Westergaard, 1958; Charlesworth, 2016). The Y chromosome, which never recombines, experiences reduced selection, which results in an accumulation of deleterious mutations and transposable elements (Charlesworth & Charlesworth, 2000). This phenomenon of Y degeneration is expected to gradually increase the size of the Y chromosome initially, and then to reduce it (Ming *et al*., 2011). Therefore, after sufficient time of divergence, we expect to observe chromosome heteromorphy, *i*.*e*. a Y chromosome larger or smaller than the X chromosome, depending on the progress of degeneration (Ming *et al*., 2011). In plants, dioecy is often of recent origin (Käfer *et al*., 2017), thus limiting the age of the sex chromosomes. Indeed, several rather recently evolved (less than 10 million years (My) old) homomorphic sex chromosome systems with small non-recombining regions have been described, as in *Carica papaya* and *Asparagus officinalis* (Wu & Moore, 2015; Harkess *et al*., 2017). Heteromorphic sex chromosome systems are also found, with the Y being larger than the X, but recombination suppression happened also relatively recently (less than 20 My ago), as in *Silene latifolia* and *Coccinia grandis* (Sousa *et al*., 2013; Krasovec *et al*., 2018; Fruchard *et al*., 2020). A few cases in which dioecy evolved longer ago also exist (Käfer *et al*., 2017), but no strongly degenerated sex chromosomes have been described so far. Pucholt *et al*. (2017) described very young sex chromosomes in *Salix viminalis* despite ancestral dioecy for the sister genera *Salix* and *Populus*. Thus, either the sex chromosomes evolved independently in different species, or there have been frequent turnovers. In the fully dioecious palm tree genus *Phoenix*, a sex-linked region evolved before the speciation of the fourteen known species (Cherif *et al*., 2016; Torres *et al*., 2018). These sex chromosomes might be old, but do not appear to be strongly differentiated. A similar situation has been reported in the grapevine (*Vitis*) genus (Badouin *et al*., 2020; Massonet *et al*., 2020), possibly because sex chromosome evolution is slowed down in such perennials with long generation time (Muyle *et al*., 2017).

Thus, to our knowledge, no sex chromosomes shared by species in different genera have been described in plants so far, a situation in stark contrast to the animals, for which several systems are more than 100 My old and are shared by whole classes, *e*.*g*. birds and mammals (Ohno, 1969; Fridolfsson *et al*., 1998; Cortez *et al*., 2014).

Dioecy very likely evolved before the genera *Cannabis* and *Humulus* split, and might even be ancestral in the Cannabaceae family (Yang *et al*., 2013; Zhang *et al*., 2018). *Cannabis sativa* (marijuana and hemp) is a dioecious species with nearly homomorphic XY chromosomes. These sex chromosomes have a large non-recombining region and are estimated to have started diverging between 12 and 28 My ago (Peil *et al*., 2003; Divashuk *et al*., 2014, Prentout *et al*., 2020). As for *C. sativa*, cytological analyses of *Humulus lupulus* (hop) found a XY chromosome system with a large non-recombining region, but the Y chromosome is smaller than the X (Shephard *et al*., 2000; Karlov *et al*., 2003; Divashuk *et al*., 2011). The divergence between *H. lupulus* and *C. sativa* is estimated between 21 and 25 My old (Divashuk *et al*., 2014; Jin *et al*., 2020), which is lower than our higher bound estimate of the age of the *C. sativa* sex chromosomes (Prentout *et al*., 2020). It is thus possible that the sex chromosomes of *C. sativa* and *H. lupulus* evolved from the same pair that already stopped recombining in their common ancestor, a question we address here.

As in many cultivated dioecious species, only female hop plants are harvested. Hop is used in beer brewing for its bitterness, and its production is increasing worldwide (Neve, 1991; King & Pavlovic, 2017), mostly because of the craft beer revolution (Barth-Haas, 2019; Mackinnon & Pavlovic, 2019). The molecule responsible for hop flowers bitterness, lupulin, is concentrated in female ripe inflorescences, called cones (Okada & Ito, 2001). In pollinated cones, the presence of seed reduces their brewing quality; since *H. lupulus* is wind pollinated, a single male plant in the hop field or its vicinity can cause broad scale damage to the crop (Thomas & Neve, 1976). Usually, hop is not grown from seeds, so female-only cultures are easy to obtain, and there is no need for large-scale early sexing as in *Cannabis sativa* (cf. Prentout *et al*., 2020). However, for varietal improvement where controlled crosses are needed, knowing the sex early might be beneficial. In *H. lupulus*, the identification of the sex is reliable 1-2 years after the sewing (Conway and Snyder, 2008; Patzak *et al*., 2002). A few markers have been developed, but the use of Y-specific coding sequences may increase the marker quality (Patzak *et al*., 2002, Cerenak *et al*., 2019).

Here we sequenced the transcriptome of fourteen *H. lupulus* individuals. These individuals came from a cross, from which we sequenced the parents and six offspring of each sex. We used the probabilistic approach SEX-DETector, which is based on allele segregation analysis within a cross, to identify sex-linked sequences (Muyle *et al*., 2016). From theses analyses on *H. lupulus* and our previous results on *C. sativa* (Prentout *et al*., 2020) we describe for the first time well-differentiated sex chromosomes shared by two different genera in plants.

## Materials and Methods

### Biological material and RNA-sequencing

As indicated in Fig. **1a**, we realized a controlled cross for sequencing. The *H. lupulus* parents, cultivar ‘Wye Target’ (WT; female) and the Slovenian male breeding line 2/1 (2/1), as well as 6 female and 6 male F1 siblings (Jakše *et al*., 2013) were collected in July 2019 in the experimental garden of Slovenian Institute of Hop Research and Brewing, Žalec.

**Figure 1.**
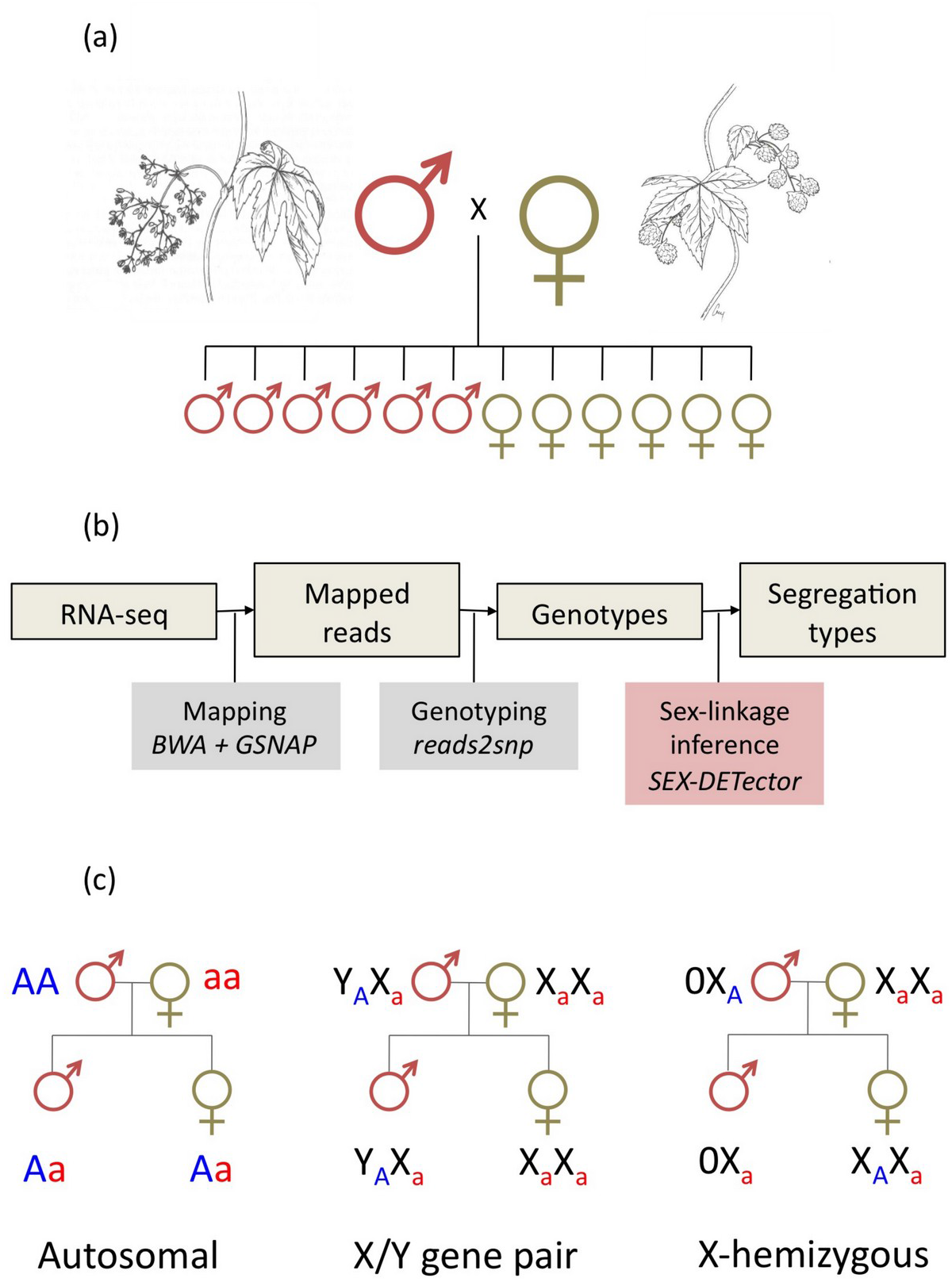
Schematic representation of the workflow used to detect sex-linkage. **(a)** Experimental design: six females and six males were obtained by a controlled cross, and all individuals (14) were sequenced. **(b)** Bioinformatic pipeline for the treatment of RNAseq data. **(c)** Illustration of the underlying principles of the SEX-DETector segregation analysis.

All offspring were phenotypically confirmed to carry either male or female reproductive organs and showed no anomalies in the microsatellite genotyping data (Jakše *et al*., 2013). Young leaves from the laterally developing shoots were picked, wrapped in aluminum foil and flash frozen *in situ* in liquid nitrogen. Later they were pulverized and stored at -80°C until RNA isolation.

Total RNA was isolated from 100 mg frozen tissue pulverized in liquid nitrogen according to the protocol of Monarch Total RNA Miniprep Kit, including removal of DNA from the column with DNase I (New England Biolabs). Total RNA was quantified with Qbit 3.0, and quality was verified with the Agilent RNA Nano 6000 Kit to confirm appropriate sample RIN numbers. The total RNA samples were sent to Novogen for mRNA sequencing using Illumina’s 100 bp paired end service. The data were submitted to the SRA database of the NCBI (BioSample accession SAMN17526021).

### Mapping, genotyping and SEX-DETector

The bioinformatic pipeline is schematically indicated in Fig. **1b**. The RNA-seq data were mapped on the transcriptome of *H. lupulus* (Padgitt-Cobb *et al*., 2019) and the transcriptome of *C. sativa* that we also used for our previous *C. sativa* analysis (Supporting Information; Van Bakel *et al*., 2011; Prentout *et al*., 2020). For the mapping, we ran GSNAP (version 2019-09-12; Wu and Nacu, 2010; Wu *et al*., 2016), an aligner that enables SNP-tolerant mapping, with 10% mismatches allowed. This approach, already used for *C. sativa* analysis, increased through several iterations the mapping quality by adding Y-specific SNPs to the references (and *H. lupulus* specific SNPs while mapping on *C. sativa* reference; see Prentout *et al*., 2020). Then, SAMTOOLS (version 1.4; Li *et al*., 2009) was used to remove unmapped reads and sort mapping outputs files for the genotyping. We genotyped individuals with reads2snp (version 2.0.64; Gayral *et al*., 2013), as recommended for SEX-DETector (Muyle *et al*., 2016), *i*.*e*., by accounting for allelic expression biases, without filtering for paralogous SNPs, and only conserving SNPs supported by at least three reads for subsequent analysis.

We ran the XY model of SEX-DETector on the genotyping data, using the SEM algorithm and a threshold for an assignment of 0.8. SEX-DETector computes a posterior probability of being autosomal (P_A_), XY (P_XY_) and X-hemizygous (P_X-hemi_) for each SNP and for each gene (Fig. **1c**). Thus, a gene with a P_A_ greater than or equal to 0.8 and at least one autosomal SNP without genotyping error is classified as ‘autosomal’; a gene with P_XY_ + P_X-hemi_ greater than or equal to 0.8 and at least one sex-linked SNP without genotyping error is classified as ‘sex-linked’; otherwise, the gene is classified as ‘lack-of-information’. Among the sex-linked genes, X-hemizygous ones are those with only X-hemizygous SNPs and at least one without genotyping error, as well as genes with a Y expression detected only on aberrant SNPs (see Muyle *et al*., 2016). A parameter that is important to optimize with SEX-DETector is the Y specific genotyping error rate (*p*; see Muyle *et al*., 2016). However, the Y mapping quality reduces with the XY divergence, therefore, old and highly divergent sex chromosomes are more susceptible to mapping errors and thus genotyping errors. *p* is expected to be close to the whole transcriptome genotyping error rate (*ε*), but could be higher due to weak expression (resulting in less reads) of the Y copies of genes. To reduce the gap between these two error rates, we ran 4 iterations with GSNAP, using at each time the SNPs file output from SEX-DETector to increase reference quality.

### Sex-linked gene positions on *C. sativa* genome

We determined the position of the transcript sequences, used for the mapping, on a chromosome-level assembly of the *C. sativa* genome (Grassa *et al*., 2018) with blast (version 2.2.30+; Altschul *et al*., 1990). We selected the best hit with an e-value lower than 10^−4^ to determine the position of the transcript on the genome. Then, we split each chromosome in windows of 2 Mb and computed the density of sex-linked genes and non-sex-linked genes per window using BEDTOOLS (version 2.26.0; Quinlan & Hall, 2010). Proportions of sex-linked genes were computed by dividing the number of sex-linked genes by the total number of genes (sex-linked, autosomal, and undetermined) in the same window. For *C. sativa*, densities were already available from our previous analysis (Prentout *et al*., 2020).

### Molecular clock and age of sex chromosomes

We used the translated reference transcripts (van Bakel *et al*., 2011) to determine the X and Y Open Reading Frame (ORF) of nucleotide reference transcripts. For each XY gene pair, the *dS* values were estimated with codeml (PAML version 4.9; Yang, 2007) in pairwise mode. Then, we used two molecular clocks, derived from *Arabidopsis* species, to estimate the age of *H. lupulus* sex chromosomes (Koch *et al*., 2000; Ossowski *et al*., 2010). In the wild, *H. lupulus* flowers in the second or third year of the development (Patzak *et al*., 2002; Polley *et al*., 1997), therefore, we took a generation time (GT) of 2 years, and used the molecular clock as follows: (GT *×dS*)/ *rate*=*dS*/ (7.5 *×*10^*−* 9^) using the molecular clock from Koch *et al*. (2000); (GT *×dS*)/ (2*×µ*)=*dS* /(7 *×*10^*−* 9^) using the clock from Ossowski *et al*. (2010). Three different estimates of *dS* were used: the maximum *dS* value, the mean of 5% highest *dS* values, and the mean of 10% highest *dS* values.

### X and Y allele-specific expression analysis

In addition to identifying X and Y alleles, SEX-DETector estimates their expression based on the number of reads (Muyle *et al*., 2016). These estimates rely on counting reads spanning XY SNPs only and were normalized using the total read number in a library for each individual. They were further normalized by the median autosomal expression for each individual. *C. sativa* results presented here were generated in our previous analysis on *C. sativa* sex chromosomes (Prentout *et al*., 2020).

### Correction of Y read mapping bias

The use of a female reference for the mapping of the reads might create mapping biases, resulting in the absence of Y reads in the mostly diverging parts of the genes. This issue may reduce the divergence detected and change the phylogenetic signal (Dixon *et al*., 2019). If, within the same gene, regions that lack Y reads coexist with regions were the Y reads correctly mapped, we expect to see a signature similar to gene conversion, *i*.*e*. region-wise variation in the divergence. Therefore, we ran geneconv (version 1.81a; Sawyer, 1999) in pairwise and group mode with the multiple alignments used for the phylogeny (on 85 gene alignments before Gblock filtering, see below) in order to identify and remove regions with reduced divergence. We defined two groups, one for X and Y sequences in *H. lupulus* and the other one for X and Y sequences in *C. sativa*. Then, we conserved only inner fragments and split the gene conversion regions from regions without gene conversion to obtain two subsets per genes. Thus, we obtained a subset of sequences corrected for the mapping bias, in addition to the set of genes not filtered with geneconv.

### Phylogenetic analysis

We reconstructed gene families for genes identified as sex-linked in both *C. sativa* and *H. lupulus*. Then, we used blastp, filtering for the best hit (with an e-value threshold fixed at 10^−4^), to find homologous sequences between *C. sativa* reference transcripts (the query sequence in blastp) (van Bakel *et al*., 2011) and 4 outgroup transcriptomes (the subject sequence in blastp): *Trema orientalis* (Cannabaceae; van Velzen *et al*., 2018), *Morus notabilis* (Moraceae; He *et al*., 2013), *Fragaria vesca ssp. vesca* (Rosaceae; Shulaev *et al*., 2011), and *Rosa chinensis* (Rosaceae; Raymond *et al*., 2018). Gene families for which at least two outgroup sequences have been identified were conserved, incomplete gene families were discard from subsequent analysis. Then, we added X and Y sequences reconstructed by SEX-DETector to each gene families. To identify potential paralogous sequences or variants from alternative splicing, a blast of all sequences *vs* all sequences was realized. If two sequences from two distinct gene families blast with each other (with an e-value threshold fixed at 10^−4^), then both families were removed from the dataset. Finally, we retrieved the corresponding nucleotidic sequences of each protein families, which constituted the dataset used for the phylogenetic analysis.

Using Macse (version 2.03; Ranwez *et al*., 2011), and before alignment, non-homologous segments of at least 60 nucleotides within or 30 nucleotides at the extremity of a nucleotide sequence were trimmed if they showed less than 30 % of dissimilarity compared to any other sequences in the gene family. This step allowed to remove misidentified outgroup sequences, then, gene families with no remaining outgroup sequences were discarded. Finally, remaining families were aligned with Macse, allowing sequences to be removed and realigned, one sequence at a time and over multiple iterations, to improve local alignment.

Nucleotide alignments were cleaned at the codon level using Gblocks (with default parameters) to conserve only codons shared by all sequences (version 0.91b; Castresana, 2000). For maximum-likelihood (ML) phylogenetic tree reconstruction, we used ModelFinder in IQ-TREE (version 1.639; Nguyen *et al*., 2015; Kalyaanamoorthy *et al*., 2017) to select the best-fit substitution model for each alignment. Those models were then used in RAxML-NG (version 1.0.0; Kozlov *et al*., 2019) to reconstruct gene family trees. The number of bootstrap replicates was estimated using autoMRE (Pattengale *et al*., 2010) criterion (maximum 2,000 bootstraps). The ML phylogenetic tree reconstruction was run on two datasets, one without removing potential mapping biases, and one with the potential mapping bias removed, as described above.

Bayesian phylogenies were built using Phylobayes (version 3.4; Lartillot *et al*., 2009) with the site-specific profiles CAT and the CAT-GTR models with a gamma distribution to handle across site rate variations. Two chains were run in parallel for a minimum of 500 cycles. The convergence between the two chains was checked every 100 cycles (with a burn-in equal to one fifth of the total length of the chains). Chains were stopped once all the discrepancies were lower or equal to 0.1 and all effective sizes were larger than 50 and used to build a majority rule consensus tree.

### Statistics and linear chromosome representations

The statistical analyses have been conducted with R (version 3.4.4; R Core Team, 2013). We report exact p-values when they are larger than 10^−5^. The representation of phylogenetic topologies, *dS* values on the first chromosome and the dosage compensation graphics have been done with ggplot2 (Wickham, 2011). For the circular representation of the sex-linked genes density along the *C. sativa* genome we used Circos (version 0.69-6; Krzywinski *et al*., 2009). We calculated confidence intervals for the median of a dataset of *n* observations by resampling 5000 times *n* values from the dataset (with replacement). The confidence intervals are then given by the quantiles of the distribution of median values obtained by resampling.

## Results

### Identification of sex-linked genes in *H. lupulus*

As detailed in the Supporting Information, we retained the *C. sativa* reference transcriptome for downstream analysis based on the quality of the mapping and the results of SEX-DETector.

Of the 30,074 genes in the *C. sativa* reference transcriptome, 21,268 had detectable expression in our *H. lupulus* transcriptome data. The difference of properly-paired mapped reads between males (mean: 32.3%) and females (mean: 34.9%) is slightly significant (Wilcoxon’s test two-sided p-value = 0.038, see Supporting Information Table S1), which may be explained by a lack of Y-specific reads mapping on the female reference.

The sex-linked sequences from *H. lupulus* transcriptome data were identified with SEX-DETector (Muyle *et al*., 2016). It is important that genotyping error rate parameters *ε and p* have similar values (*ε*: whole transcriptome; *p*: Y chromosome) to obtain reliable SEX-DETector outputs. At the fourth iteration of GSNAP mapping on *C. sativa* reference transcriptome *ε* and *p* stabilized at 0.06 and 0.20, respectively (Supporting Information Table S2). Upon closer inspection, one *H. lupulus* male (#3) appeared to have many genotyping errors, as for some XY genes, this male was genotyped both heterozygous (XY) and homozygous (XX), which increased the error rate *p*. A particularly strong Y reads mapping bias in this male may explain these observations. After removal of this male, the error rate *p* dropped to 0.10 (Supporting Information Table S2). A total of 265 sex-linked genes were identified in *H. lupulus*, which represents 7.8% of all assigned genes (Table 1).

**Table 1.**
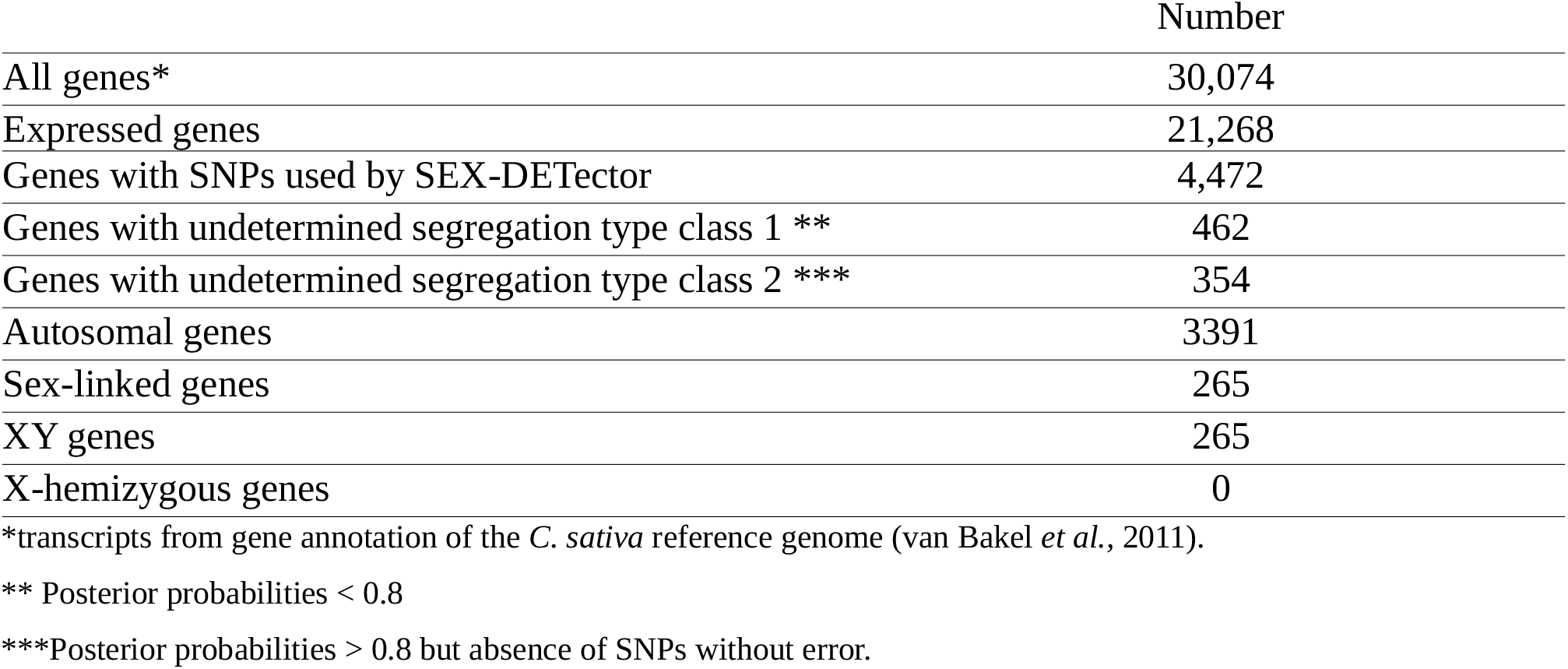
Summary of the SEX-DETector results.

|  | Number |
| --- | --- |
| All genes* | 30,074 |
| Expressed genes | 21,268 |
| Genes with SNPs used by SEX-DETECTOR | 4,472 |
| Genes with undetermined segregation type class 1 ** | 462 |
| Genes with undetermined segregation type class 2 *** | 354 |
| Autosomal genes | 3391 |
| Sex-linked genes | 265 |
| XY genes | 265 |
| X-hemizygous genes | 0 |

\*transcripts from gene annotation of the *C. sativa* reference genome (van Bakel *et al.*, 2011).
\*\* Posterior probabilities < 0.8
\*\*\*Posterior probabilities > 0.8 but absence of SNPs without error.

### *H. lupulus* and *C. sativa* sex chromosomes are homologous

Among 265 *H. lupulus* XY genes from the *C. sativa* transcriptome assembly (van Bakel *et al*., 2011), 254 genes are present on the *C. sativa* chromosome-level genome assembly (Grassa *et al*., 2018). As shown in Figure 2, 192 of these genes (75.6%) map to on *C. sativa* chromosome number 1, which is the chromosome we previously identified as the X chromosome in *C. sativa* (Prentout *et al*., 2020). Of the 265 sex-linked genes in *H. lupulus*, 112 were also detected as sex-linked in *C. sativa*, while 64 were detected as autosomal and 89 had unassigned segregation type (Prentout *et al*., 2020).

**Figure 2.**
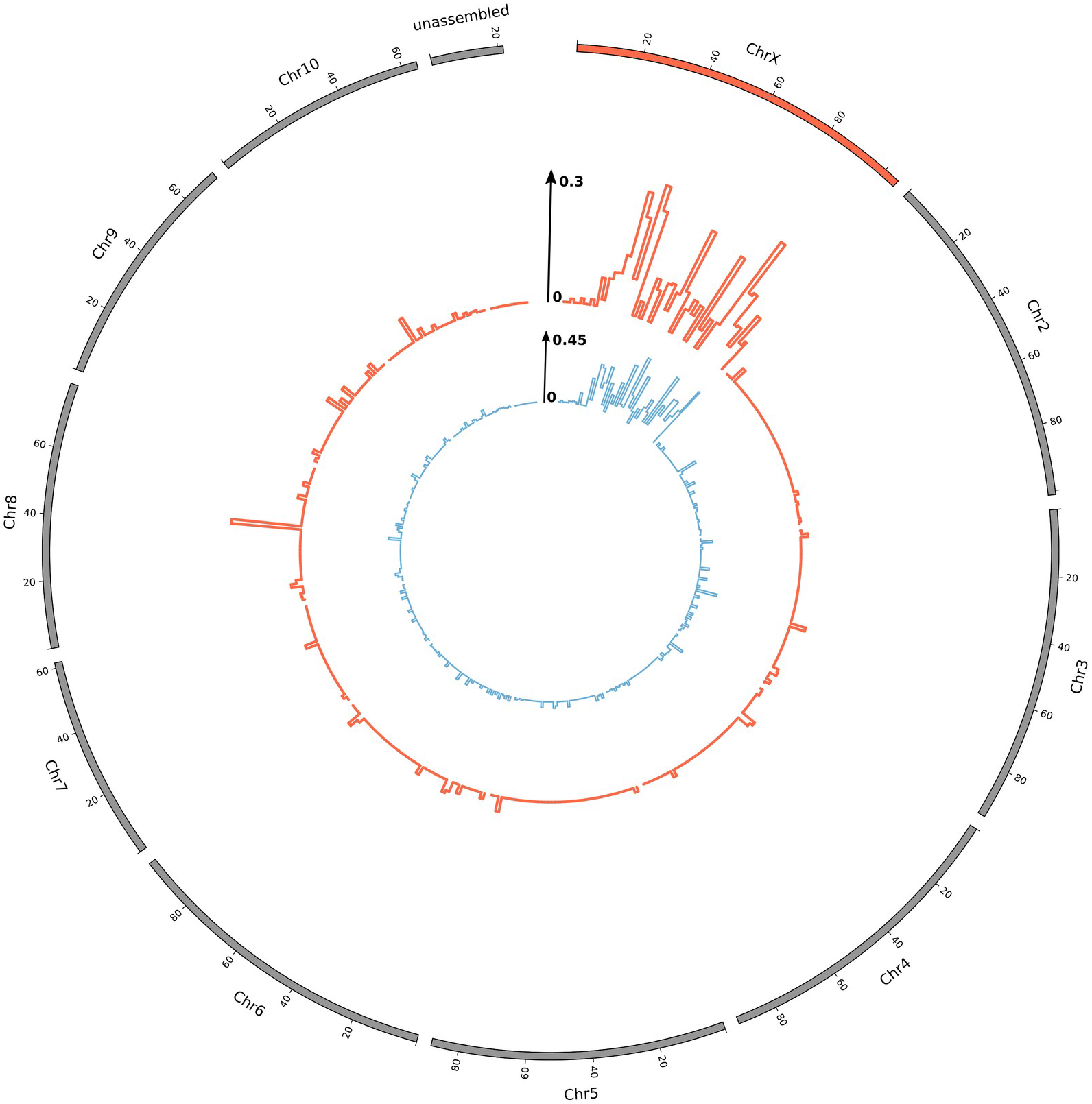
*H. lupulus* sex-linked genes mapped on the *C. sativa* genome (Grassa *et al*., 2018). Inner graphs (in blue): *C. sativa* sex-linked gene density corrected by the total gene density in 2-Mb windows (from Prentout *et al*., 2020). Outer graphs (in red): *H. lupulus* sex-linked gene density corrected by the total gene density in 2-Mb windows. Chromosome positions are given in Megabases.

The synonymous divergence (*dS*) between X and Y copies of the sex-linked genes of *H. lupulus* is distributed similarly along the *C. sativa* sex chromosome as the values for this latter species, as shown in Figure 3. While the sampling variation of these *dS* values is large, as expected, it can be observed that these values tend to be larger in the region above 75 Mb.

**Figure 3.**
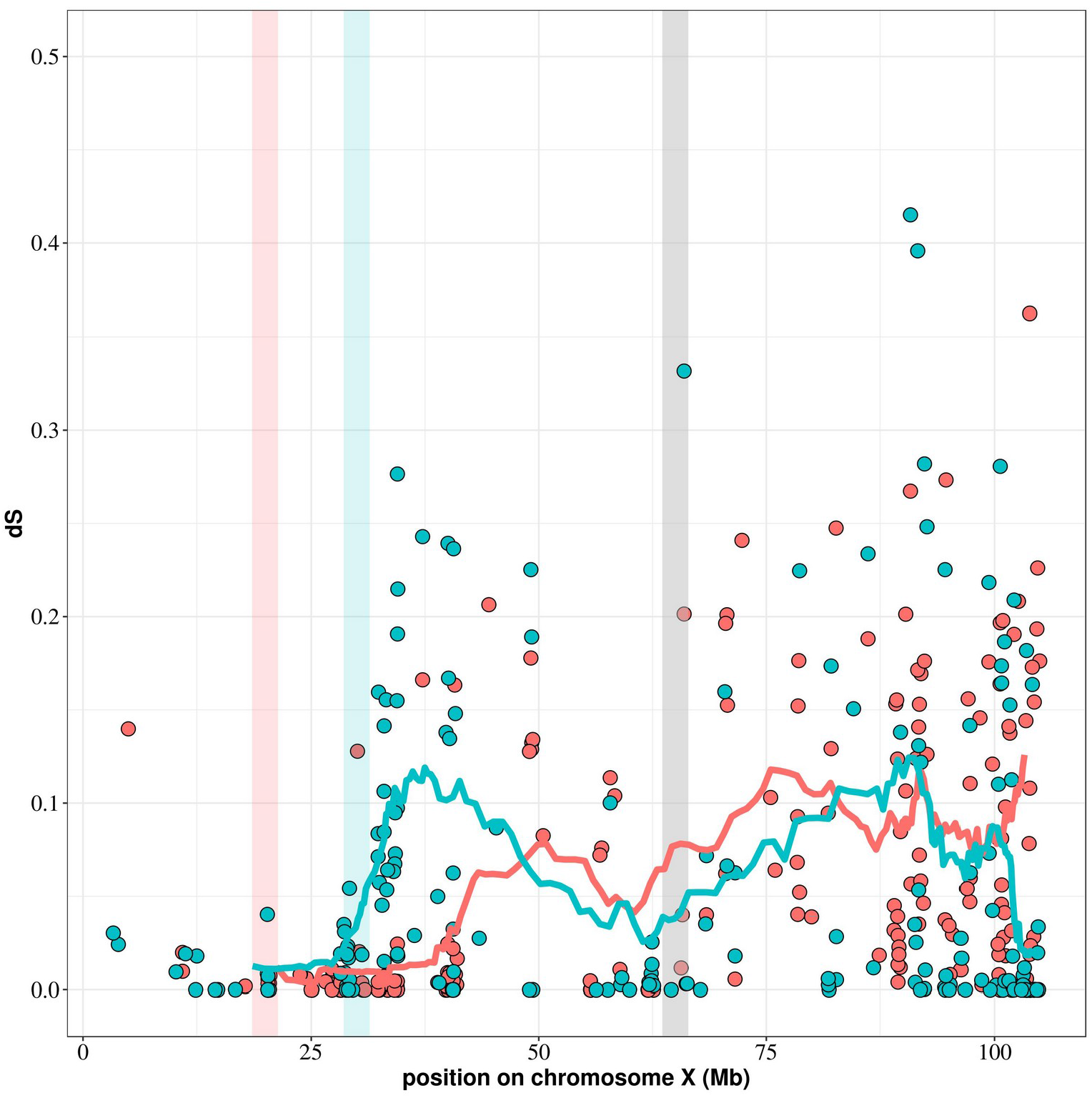
Synonymous divergence (*dS*) between X and Y copies of *H. lupulus* sex-linked genes (red) and those of *C. sativa* (blue) along the X chromosome of *C. sativa*. The curves represent the *dS* with sliding windows (windows of 20 points), for *H. lupulus* (red) and *C. sativa* (blue). The vertical red bar represents the putative Pseudo-Autosomal Boundary (PAB) in *H. lupulus*, the vertical blue bar represents the putative PAB in *C. sativa*, the vertical grey bar represents the putative boundary between the region that stopped recombining in the common ancestor and the region that stopped recombining independently in the two species (see below).

### X-Y recombination stopped before *Cannabis* and *Humulus* genera split

We reconstructed phylogenetic trees of genes detected as sex-linked in both species, including outgroup sequences from the order Rosales. For 27 out of the 112 sex-linked genes present in both species, we could not identify any homolog in the outgroup species and those genes were excluded from further analysis. For the remaining 85, we quantified the topology of the gametologous sequences in the Cannabaceae, considering a node as well resolved when the bootstrap support exceeded 95%, or Bayesian support exceeded 0.95.

The three different methods for phylogenetic reconstruction provided consistent phylogenies (Table 2). More precisely, we observed three major topologies, as shown in Figure 4: X copies of both species form a clade separated from a clade of Y sequences (topology I, Fig. **4a**), the X and Y sequences of each species group together (topology II, Fig. **4b**), or a paraphyletic placement of the X and Y sequences of *H. lupulus*, relative to *C. sativa* sequences (topology III, Fig. **4c**). As shown in Table 2, we found that most genes had topology II, corresponding to recombination suppression after the split between the species. A few genes, however, had topology I, which corresponds to genes for which recombination was suppressed in a common ancestor of both species. As shown in Fig. **4d**, topologies I and III occurred mainly beyond 80 Mb, while topology II occurred all over the chromosome. Topology I is associated with higher synonymous divergence.

**Table 2.**
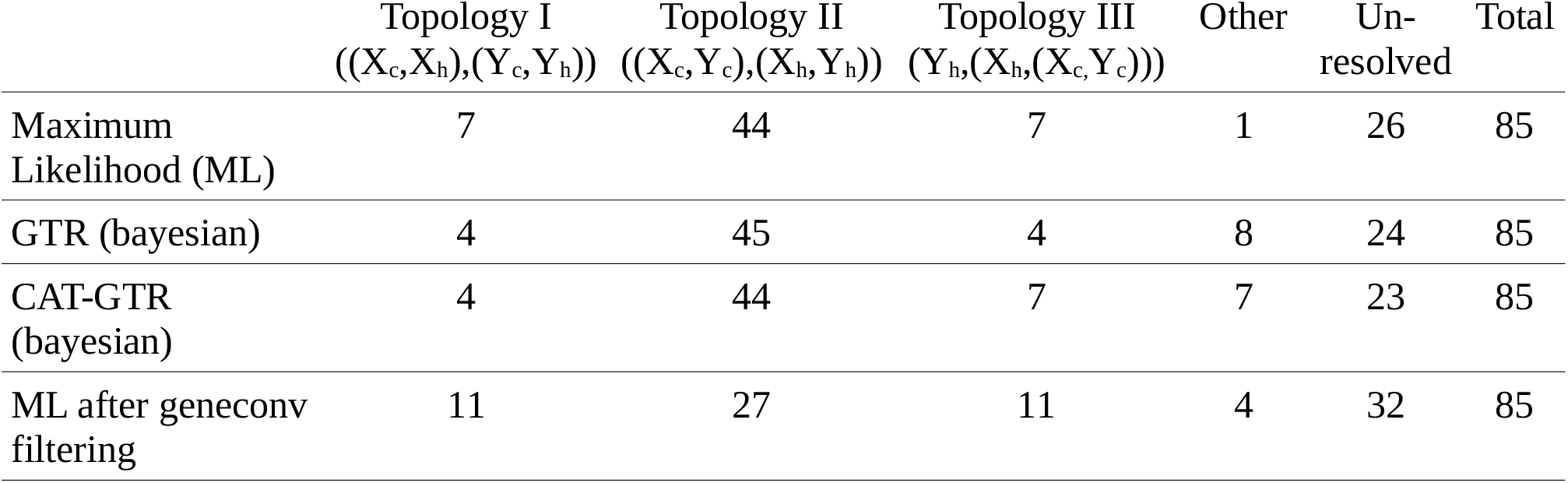
Results of the phylogenetic reconstruction of sex-linked genes. Phylogenetic trees with a bootstrap equal or greater than 95% (and posterior probabilities higher than 0.95 for Bayesian reconstructions) at the node separating *C. sativa* and *H. lupulus*, or Y and X sequences, are presented in the first four columns. Phylogenetic trees without such support are classified as ‘unresolved’.

**Figure 4.**
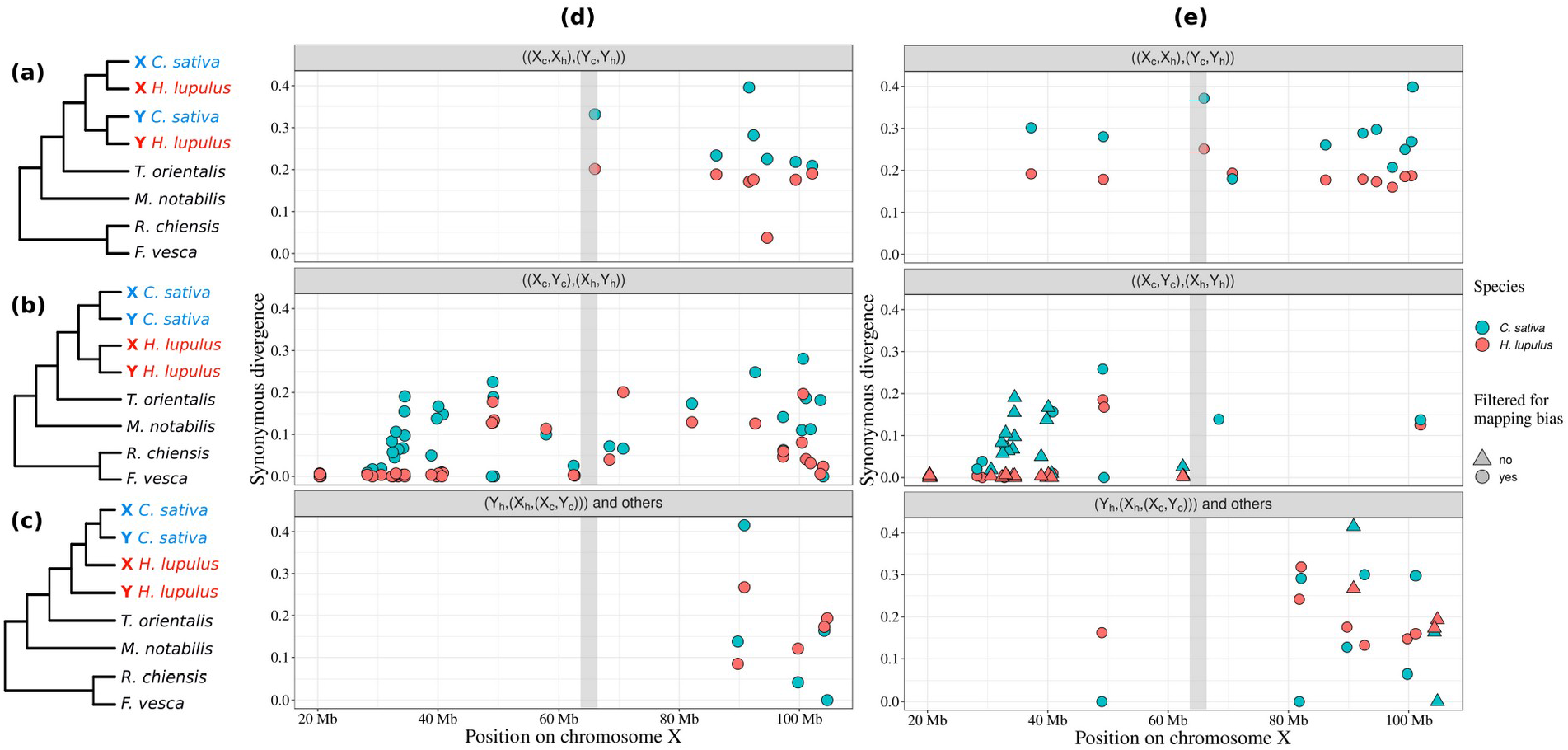
Distribution of the three topologies of the sex-linked genes on the X chromosome: **(a)** Topology I, XX-YY – arrest of recombination older than the split between the two genera, **(b)** Topology II, XY-XY – arrest of recombination younger than the split between the two genera, **(c)** Topology III, Y-X-XY – *H. lupulus* X chromosome is closer to *C. sativa* sequences than its Y counterpart. **(d)** Distribution of the topologies along the *C. sativa* X chromosome (‘other’ topology is included in the Y-X-XY topology panel), using the full gene sequences. For each gene, dots represent the *dS* values in *C. sativa* (blue) and *H. lupulus* (red). **(e)** Distribution of the topologies after filtering out possible mapping biases through geneconv. Triangles indicate that at least one segment was removed, dots indicate sequences for which no mapping bias was detected.

We identified 42 genes, out of the 85 genes used for the phylogeny, with at least one fragment in at least one species that displayed reduced divergence (with a p-value < 0.05 in geneconv output). Because this reduction of divergence may be caused by a mapping bias of Y reads, we ran the ML phylogenetic reconstruction method on regions with and without mapping bias (example in Supporting Information Fig. S7). As shown in Table 2 and Fig. **4e**, representing genes filtered for mapping bias, the proportion of genes displaying topology I, indicating recombination suppression in the most recent common ancestor, increased, while less genes with topology II were mainly found in a restricted region corresponding to the region where recombination stopped independently between the two species.

The vertical grey bar (panels **(d)** and **(e)**) represents the putative boundary between the region that stopped recombining in the common ancestor and the region that stopped recombining independently in the two species (see below).

This leads us to define three regions on the X chromosomes of *C. sativa* and *H. lupulus* (with the *C. sativa* X chromosome as a reference). A region from ∼65Mb to the end of the X chromosome that stopped recombining in the last common ancestor; from ∼20-30Mb to ∼65Mb, a part of the non-recombining region that evolved independently in the two species; and from the beginning of the chromosome to ∼20-30Mb, the pseudo-autosomal region (PAR), where few sex-linked genes are found.

### Age of *H. lupulus* sex chromosomes

To estimate the age of the sex chromosomes, we used the maximum synonymous divergence between X and Y sequences and two molecular clocks, which were both derived from *Arabidopsis*. Because the sampling variance in *dS* values can be large, we used three ways to calculate the maximum *dS* value: the single highest *dS* value; the average of the 5% highest values; and the average of the 10% highest values. Furthermore, we calculated these on the raw alignments as well as the alignments with possible mapping biases removed. The different estimates, all calculated assuming a generation time of 2 years, are given in Table 3, and yield values between 29 and 51.4 My. Minimal synonymous divergence between *C. sativa* and outgroup species *Morus notabilis* and *Rosa chinensis* is ∼0.45 and ∼0.65, respectively (Supporting Information Fig. S5 and Fig. S6), higher than the maximum synonymous divergence between sex-linked gene copies, indicating that the sex chromosomes probably evolved in the Cannabaceae family.

**Table 3.** Age estimates (in millions of years, My) with two molecular clocks and different maximum *dS* values, for a generation time of two years. For each estimation of the *dS* value, two ages were obtained using the molecular clocks of ^1^ Ossowski *et al*. (2010) and ^2^ Koch *et al*. (2000). Two alignment datasets were used, with or without filtering for possible mapping bias.

|  | No filtering |  |  | Mapping bias filtering |  |  |
| --- | --- | --- | --- | --- | --- | --- |
|  | *dS* | age (My)<sup>1</sup> | age (My)<sup>2</sup> | *dS* | age (My)<sup>1</sup> | age (My)<sup>2</sup> |
| Highest *dS* | 0.362 | 51.4 | 48.0 | 0.362 | 51.4 | 48.0 |
| Mean top 5% | 0.249 | 35.6 | 33.2 | 0.274 | 39.1 | 36.5 |
| Mean top 10% | 0.217 | 31.0 | 29.0 | 0.214 | 34.4 | 32.1 |

### Y gene expression

The Y over X expression ratio is a standard proxy for studying the degeneration of the Y chromosome. An Y/X expression ratio close to 1 means no degeneration, an Y/X expression ratio close to 0.5 or below means strong degeneration. In *H. lupulus*, the median Y/X expression ratio is equal to 0.637 (Supporting Information Fig. S1), which is significantly different from 1 (99^th^ percentile of median distribution with 5,000 samples in initial distribution = 0.673, see methods). The median is not different when considering all sex-linked genes (0.637) or only the sex-linked genes mapping on *C. sativa* X chromosome (0.639, p-value = 0.70, one-sided Wilcoxon rank sum test).

In both species, the reduced Y expression is correlated to the position on the X chromosome (linear regression: adjusted *R^2^* = 0.134, p-value < 10^−5^; and adjusted *R^2^* = 0.278, p-value < 10^−5^ for *H. lupulus* and *C. sativa*, respectively). As shown is Figure 5, the Y/X expression ratio is decreasing while moving away from the PAR in *H. lupulus*, and this is also confirmed in *C. sativa*.

**Figure 5.**
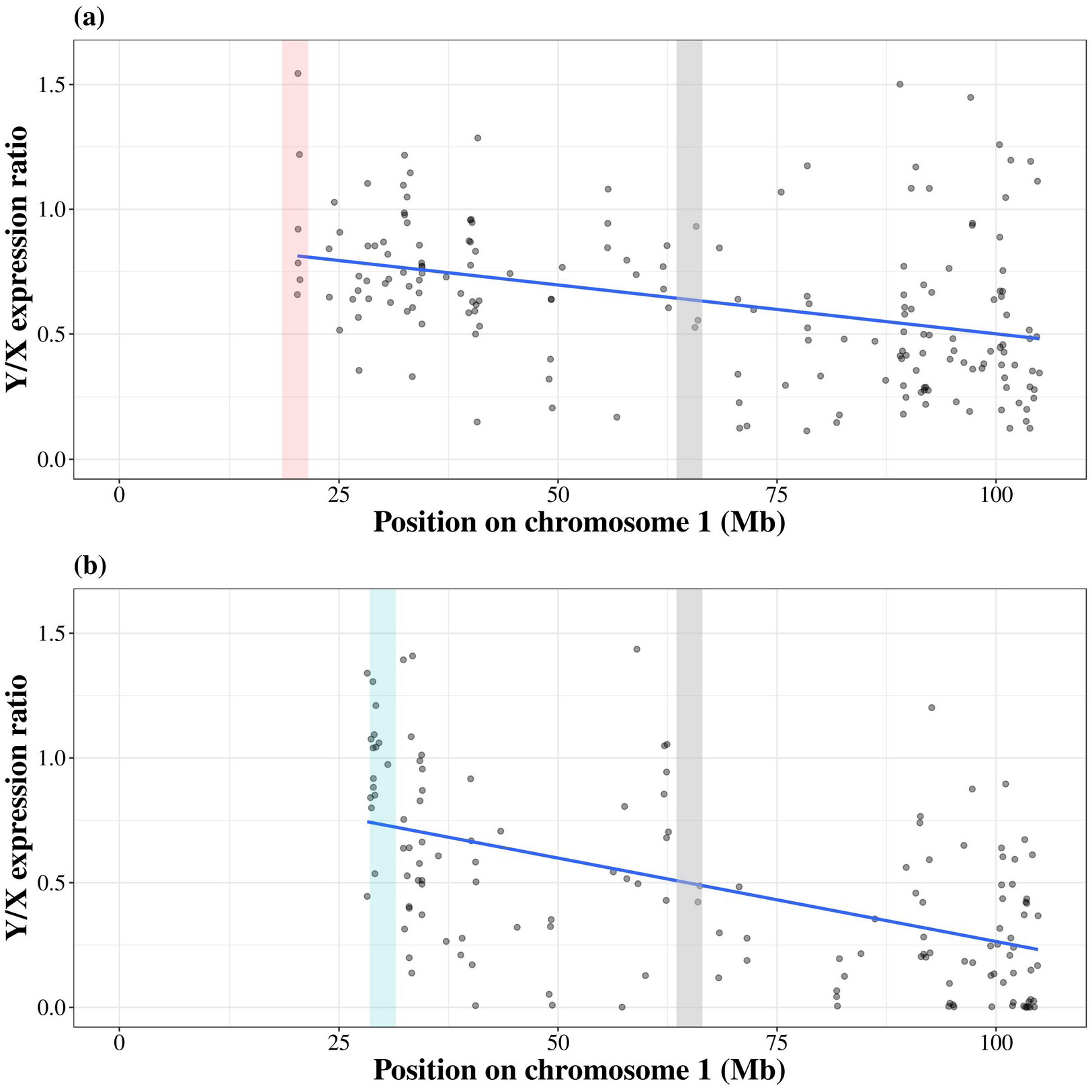
Y/X expression ratio along the X chromosome in *H. lupulus* (a), and *C. sativa* (b). The grey dots represent the Y/X expression ratio for each gene in the non-recombining region only. The blue line represents a linear regression. The vertical red bar represents the putative PAB in *H. lupulus*, the vertical blue bar represents the putative PAB in *C. sativa*, the vertical grey bar represents the putative boundary between the region that stopped recombining in the common ancestor and the region that stopped recombining independently in the two species.

#### 3 Dosage compensation

We tested whether the expression of the X chromosome has changed following degeneration of the Y chromosome, a phenomenon called dosage compensation (Muyle *et al*., 2017). We used the ratio of the male X expression over the female XX expression as a proxy for dosage compensation (Muyle *et al*., 2012) and Y/X expression ratio as a proxy for Y degeneration. Genes with strong degeneration (Y/X expression ratio close to zero) display an increased expression of the X in male (linear regression: adjusted *R^2^*=0.179, p-value < 10^−5^ and adjusted *R*^*2*^=0.097, p-value < 10^−5^ for *H. lupulus* and *C. sativa* respectively), as shown in Figure 6. A dosage compensation pattern was found in both in *H. lupulus* and *C. sativa* in agreement with previous work (Prentout *et al*., 2020).

**Figure 6.**
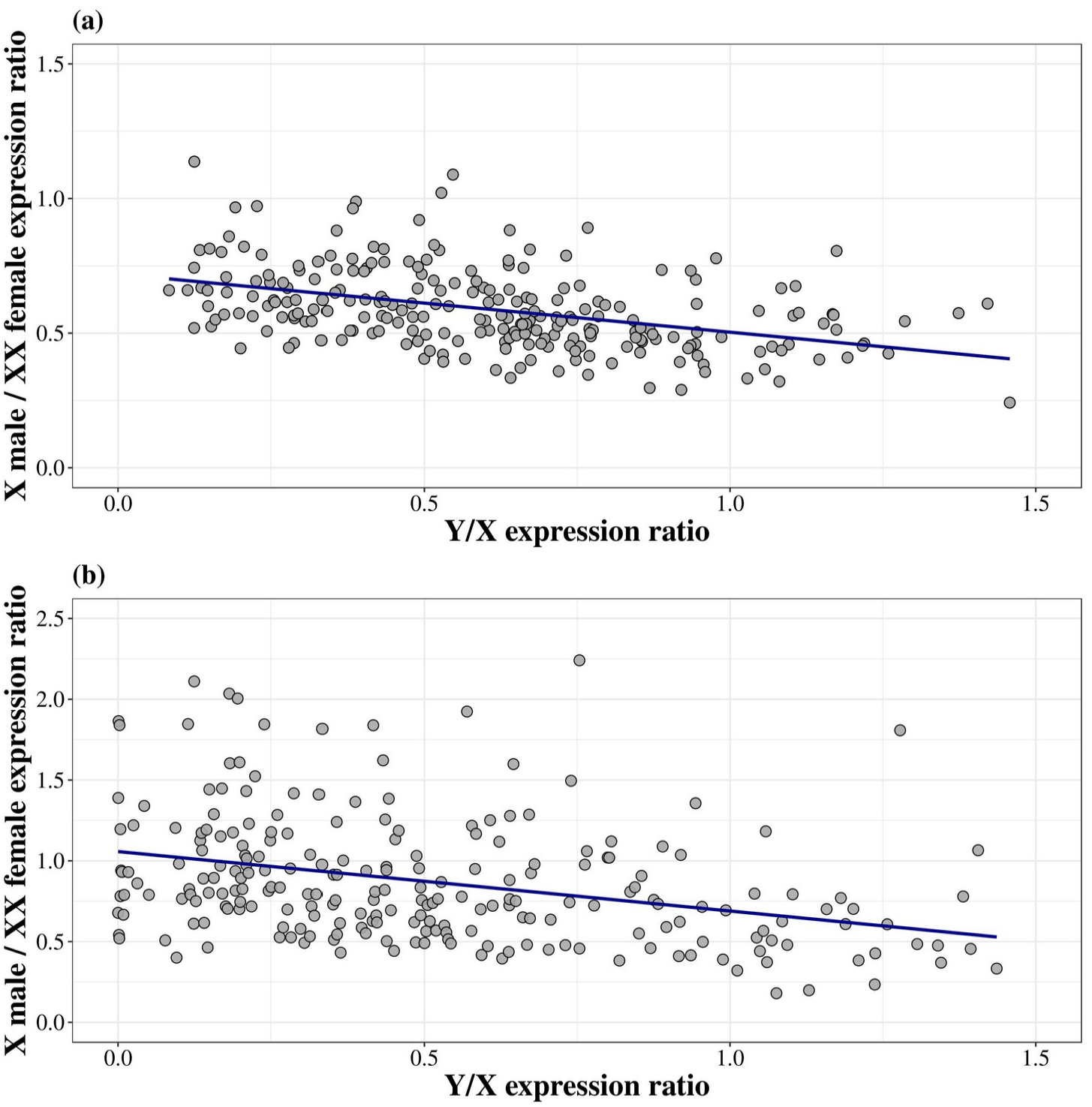
The male X expression over female XX expression versus Y/X expression ratio for *H. lupulus* (a) and *C. sativa* (b). Each black dot represents one gene. The blue line represents a linear regression.

## Discussion

We here identify the *H. lupulus* sex chromosomes, and find that they are homologous to those of *C. sativa* (Prentout *et al*., 2020), and that a part of these chromosomes had already stopped recombining in the common ancestor of the two species. Performing a segregation analysis with SEX-DETector (Muyle *et al*., 2016), we identified 265 XY genes in *H. lupulus*, among which 112 are sex-linked in *C. sativa*. Mapping these genes on the chromosome-level assembly of *C. sativa* (Grassa *et al*., 2018) suggests that the non-recombining region is large in *H. lupulus*, as proposed before, based on cytological studies (Divashuk *et al*., 2011).

We identify three different regions on the sex chromosome, based on the distribution of sex-linked gene topologies and synonymous divergence between the X and Y copies on the *C. sativa* X chromosome: one region that had already stopped recombining in the most recent common ancestor of *C. sativa* and *H. lupulus*, a region that independently stopped recombining in both species, and the pseudo-autosomal region. Our results suggest the pseudo-autosomal boundary (PAB) in *H. lupulus* may be located around position 20Mb, whereas we estimated a PAB’s position around 30Mb in *C. sativa* (Prentout *et al*., 2020); the non-recombining region may thus be larger in *H. lupulus* than in *C. sativa*. However, a chromosome-level assembly of the *H. lupulus* genome would be needed to localize the exact position of the PAB in this species, as synteny might not be fully conserved.

Several sex-linked genes had topologies that where not compatible with either recombination suppression in the most recent common ancestor or in each of the species independently. Strikingly, most of these topologies placed the *H. lupulus* Y sequence as an outgroup to the other sex-linked gene sequences. Whether this is the result of errors (*e*.*g*. long branch attraction, mapping biases) remains to be investigated. Our approach to correct for the Y read mapping read relies on geneconv, which is known to have a high rate of false negatives (Lawson & Zhang, 2009). This could also explain the unexpected presence of some of the XY-XY genes in the older region.

While recombination suppression clearly did not occur for all the sex-linked genes at the same time, we cannot distinguish whether this recombination suppression occurred gradually or stepwise, creating evolutionary strata (Charlesworth *et al*., 2005; Bergero & Charlesworth 2009; Muyle *et al*., 2017). It is unlikely this question can be addressed using synonymous divergence alone, given the important sampling variation present in this statistic. Y chromosome assemblies for both *H. lupulus* and *C. sativa* may help address this question in the future by identifying possible chromosomal inversions with respect to the X chromosomes.

We did not find X-hemizygous genes in *H. lupulus*. This is striking as 218 X-hemizygous genes (38% of all sex-linked genes) were found in *C. sativa* using the same methodology (Prentout *et al*., 2020). A very low level of polymorphism could result in the inability of SEX-DETector to identify X-hemizygous genes (Muyle *et al*., 2016), but in that case SEX-DETector should also have problems identifying autosomal genes, which was not the case here. Non-random X-inactivation in females could be an explanation, as the expression of a single X allele in females would impede SEX-DETector to identify X-linkage and X-hemizygous genes (Muyle *et al*., 2016). We ran an Allele-Specific Expression (ASE) analysis, which doesn’t support this hypothesis (Supporting Information Fig. S2, Fig. S3, Fig. S4). *H. lupulus* is probably an ancient polyploid that reverted to the ancestral karyotype (Padgitt-Cobb *et al*., 2019). It is possible however, that the *H. lupulus* X chromosome is made of two copies of the ancestral X as some cytological data seem to suggest (Divashuk et al., 2011). In this case, SEX-DETector would manage to identify the XY gene pairs, but would fail to identify the X-hemizygous genes as these genes would exhibit unexpected allele transmission patterns (Supporting Information Fig. S10).

*H. lupulus* is a rare case of XY systems in plants in which the Y is smaller than the X (cf Ming et *al*., 2011). In *C. sativa*, both sex chromosomes have similar size (Divashuk *et al*., 2014). If the size difference is caused by deletions of parts of the *H. lupulus* Y chromosome, which is the hypothesized mechanism in many species (cf Ming *et al*., 2011), we expect to observe that many XY gene pairs in *C. sativa* have missing Y copies in *H. lupulus*. As explained above, we did not detect any X-hemizygous genes. However, the XY gene pairs of *H. lupulus* are distributed uniformly on the *C. sativa* X chromosome. No region appeared to be depleted in XY genes, which is not what we should have observed if large deletions were present on the *H. lupulus* Y chromosome. The sex chromosome size differences observed in *H. lupulus* probably reflect complex dynamics, different from that of old animal systems with tiny Y chromosome due to large deletions (*e*.*g*. Skaletsky *et al*., 2003; Ross *et al*., 2005). The large size of the X chromosome in *H. lupulus* may be due to a full-chromosome duplication followed by a fusion (see above), whereas the Y chromosome has remained unrearranged. Assemblies of the *H. lupulus* sex chromosomes will be needed to test these hypotheses.

Our estimates of the age of the *H. lupulus* sex chromosomes are larger than the estimates for *C. sativa*, although we found very similar X-Y maximum divergence in both species (higher bound age estimates are ∼50My and ∼28My; highest *dS* values are 0.362 and 0.415 in *H. lupulus* and *C. sativa* respectively, see Prentout *et al*., 2020). Of course, the molecular clocks that we used are known to provide very rough estimates as they derive from the relatively distant *Arabidopsis* genus and are sensitive to potential differences in mutation rates between the annual *C. sativa* and the perennial *H. lupulus* (Neve, 1991; Petit & Hampe, 2006; Small, 2015; but see Krasovec *et al*., 2018). The difference found here mainly comes from the generation time (two years versus one year in *H. lupulus* and *C. sativa*, respectively). The short generation time in *C. sativa* is probably a derived trait, not reflecting the long-term generation time of the *Cannabis-Humulus* lineage, as the *Cannabis* genus is the only herbaceous genus in the Cannabaceae family (Yang *et al*., 2013). Thus, the remarkable similarity between the highest *dS* values in both species rather indicates that *C. sativa* and *H. lupulus* sex chromosomes have similar age, as expected if they derive from the same common ancestor, and the age estimate for *H. lupulus* recombination suppression might be the more representative one. We thus confirm here that the XY system shared by *C. sativa* and *H. lupulus* is among the oldest plant system sex chromosome documented so far (Prentout *et al*., 2020).

Dioecy was inferred as the ancestral sexual system for the Cannabaceae, Urticaceae and Moraceae (Zhang *et al*., 2018; note however that many hermaphrodite Cannabaceae were not included). We found that the synonymous divergence between the Cannabaceae species and *Morus notabilis* was about 0.45, higher than the maximum divergence of the X and Y copies in the Cannabaceae. It remains possible that the sex chromosomes evolved before the split of the Cannabaceae and Moraceae families, because the oldest genes might have been lost or were not detected in our transcriptome data. There is however no report of whether or not sex chromosomes exist in Urticaceae and Moraceae (Ming *et al*., 2011).

To estimate the Y expression, we counted the number of reads with Y SNPs. Therefore, the impact of a potential Y reads mapping bias should be weaker on Y expression analysis than on X-Y divergence analysis. We validated this assumption by removing genes with detected mapping bias from the analysis, which didn’t change the signal of Y expression reduction and dosage compensation (Supporting Information Fig. S8, Fig. S9). Dosage compensation is a well-known phenomenon in animals (*e*.*g*. Gu & Walters, 2017). It has only been documented quite recently in plants (reviewed in Muyle *et al*., 2017). Here we found evidence for dosage compensation in *H. lupulus*. This is not surprising as previous work reported dosage compensation in *C. sativa* and we showed here that both systems are homologous. *C. sativa* and *H. lupulus* add up to the list of plant sex chromosome systems with dosage compensation (see Muyle *et al*., 2017 for a review and Prentout *et al*., 2020; Fruchard *et al*., 2020 for the latest reports of dosage compensation in plants).

*H. lupulus* sex chromosomes, as those of *C. sativa*, are well-differentiated, with a large non-recombining region. Both species show similar patterns of Y degeneration and dosage compensation, despite the fact that a large part of the non-recombining region evolved independently in both species. These similarities, as well as the age of the chromosomes and the fact that they have been conserved since the most recent common ancestor of the two genera, a unique situation in plants so far, provide an exciting opportunity to test and elaborate hypotheses on sex chromosome evolution in plants.

## Acknowledgments

We thank Roberto Bacilieri for his help in setting up this collaboration and for discussions, Aline Muyle for advice on SEX-DETector and Florian Bénitière for helpful suggestions regarding graphical representations. This work was performed using the computing facilities of the CC LBBE/ PRABI; we thank Bruno Spataro and Stéphane Delmotte for cluster maintenance. Virtual machines from the Institut Français de Bioinformatique were also used to perform this work. This work received financial support from P4-0077 grant by ARRS (Slovenian Research Agency) to JJ.

## Author Contribution

Conceptualization of the study: G.A.B.M., J.K. and D.P.; methodology: G.A.B.M., J.K. N.S. and J.J.; software: D.P., T.T. and C.B.A.; formal analysis: D.P., T.T. and C.B.A.; investigation: D.P.,N.S., T.T., C.B.A., J.J., J.K., and G.A.B.M.; resources: A.C., N.S. and J.J.; writing—original draft: D.P., G.A.B.M., J.K. and T.T.; writing—review and editing: all authors; visualization: D.P. and T.T.; supervision: G.A.B.M., J.K.; project administration: G.A.B.M.; funding acquisition: N.S. and J.J.

## Data Availability

The sequence data were deposited under the Bioproject with accession number PRJNA694508, BioSample SAMN17526021 (SRR13528971; SRR13528970; SRR13528969; SRR13528968; SRR13528966; SRR13528965; SRR13528964; SRR13528967; SRR13528963; SRR13528962; SRR13528961; SRR13528960; SRR13528959; SRR13528958)

• **Regular research articles**

• **Preprint repository**

• **Web Document:**

## Supporting information legends

Table S1.

Statistics of mapping on *H. lupulus* and *C. sativa* references.

Table S2.

Summary of SEX-DETector genotyping errors and inferences

Table S3.

Expression analysis statistics summary.

Figure S1.

Histogram of the Y/X expression ratio.

Figure S2.

Histogram of the Allele-specific expression analysis for the parents.

Figure S3.

Histogram of the Allele-specific expression analysis for the daughters.

Figure S4.

Histogram of the Allele-specific expression analysis for the sons.

Figure S5.

Histogram of synonymous divergence *(dS)* between *C. sativa* and *M. notabilis*.

Figure S6.

Histogram of synonymous divergence *(dS)* between *C. sativa* and *R. chinensis*.

Figure S7.

Example of genes which topology changed with the mapping bias filtering.

Figure S8.

Y/X expression ratio along the sex chromosome without genes with a detected mapping bias.

Figure S9.

Dosage compensation analysis without genes with a detected mapping bias.

Figure S10.

SEX-DETector inference errors due to Whole Genome Duplication in *H. lupulus*.

