## Supplemental Data 1 for "Plant genera *Cannabis* and *Humulus* share the same pair of well-differentiated sex chromosomes"

### <u>Supporting Information</u>

#### <u>Materials and methods</u>

**RNA-seq mapping on assemblies, genotyping and SEX-DETector analyses**

The RNA-seq data were mapped on two references: the *Humulus lupulus* transcriptome (obtained from the genome annotation; Padgitt-Cobb *et al.,* 2019), and the *Cannabis sativa* draft transcriptome (van Bakel *et al.,* 2011). For each reference we used GSNAP with the parameters described in the main text, except that we allowed 5% of mismatches for mapping on *H. lupulus* reference instead of 10% for mapping on *C. sativa* reference. Genotyping and SEX-DETector analyses were performed in identical ways (as described in the main text).

**Synonymous divergence with outgroup species**

Synonymous divergence was estimated between *C. sativa* and *Rosa chinensis*, and *C. sativa* and *Morus notabilis*. We computed the *dS* with homologous sequences used for phylogenetic analyses (n = 85). The Open Reading Frame (ORF) was determined during the alignment step of the phylogenetic pipeline since we used protein coding sequences as guide for the nucleotide alignment. We computed *dS* with codeml as described in the main text.

**Allele-specific Expression (ASE) analysis**

We computed the ASE by counting the level of expression for each SNP with GATK (McKenna *et al.,* 2010). We retained SNPs present in at least 2 offspring and with at least 10 reads. Then, we computed the mean expression of both alleles for each genes with at least 10 SNPs. Finally, we computed the ratio: allele 1 expression / allele 2 expression.

### Results

##### Mapping results

Table S1. Mapping results on several references at fourth iteration of GSNAP. Indicated are the library sizes, and the total numbers and percentages of properly paired reads.

| Assembly | Father | Male 1 | Male 2 | Male 3 | Male 4 | Male 5 | Male 6 | Mother | Female 1 | Female 2 | Female 3 | Female 4 | Female 5 | Female 6 |
|---|---|---|---|---|---|---|---|---|---|---|---|---|---|---|
| Libraries size (in million reads) | 62.6 | 84.2 | 85.1 | 95.7 | 94.6 | 69.6 | 62.5 | 64.8 | 61.6 | 92.7 | 86.0 | 70.0 | 83.2 | 74.7 |
| *H. lupulus* transcriptome (in million reads) | 48.5 | 63.5 | 61.6 | 67.7 | 49 | 46.7 | 46.4 | 44.9 | 46.3 | 68.0 | 66.1 | 51.7 | 59.4 | 56.6 |
| *H. lupulus* transcriptome (%) | 77.3 | 75.4 | 72.3 | 70.7 | 75.9 | 67 | 74.3 | 69.3 | 75.4 | 73.4 | 76.9 | 73.8 | 71.3 | 73.8 |
| *C. sativa* transcriptome (in million reads) | 22.0 | 28.5 | 27.2 | 29.8 | 20.2 | 20.3 | 20.6 | 20.7 | 21.6 | 32.0 | 31.1 | 26.8 | 27.8 | 26.4 |
| *C. sativa* transcriptome (%) | 35.1 | 33.8 | 32.0 | 31.2 | 31.2 | 29.8 | 33.0 | 31.9 | 35.0 | 34.5 | 36.2 | 38.2 | 33.6 | 35.4 |

##### SEX-DETector outputs

Table S2. SEX-DETector genotyping error and SEX-DETector inferences, for mapping on *C. sativa* transcriptome (30,074 genes) and *H. lupulus* transcriptome (26,149 genes) references, at first and fourth GSNAP iterations. Both genotyping error rates are presented : $\varepsilon$ – Genotyping error rate for whole transcriptome; $p$ – Y genotyping error rate. Numbers of autosomal genes, XY genes and X-hemizygous genes are presented for each mapping.

| | **E** | **P** | **#Autosomal genes** | **#XY genes** | **#X-hemizygous genes** |
|---|---|---|---|---|---|
| *H. lupulus* genome 1st iteration | 0.05 | 0.88 | 970 | 0 | 0 |
| *H. lupulus* genome 4th iteration | 0.05 | 0.85 | 998 | 0 | 0 |
| *C. sativa* 1st iteration | 0.05 | 0.86 | 2948 | 0 | 0 |
| *C. sativa* 4th iteration | 0.06 | 0.20 | 3103 | 122 | 0 |
| *C. sativa* 4th iteration minus male 3 | 0.06 | 0.1 | 3391 | 265 | 0 |

##### Choice of the assembly

As explained in the main text, the optimization of the parameter $p$ is crucial to correctly interpret SEX-DETector outputs. The Table S2 shows that GSNAP mapping on *C. sativa* transcriptome was the best way to reduce $p$ to a reasonable value. Moreover, SEX-DETector didn't identify sex-linked genes with the mapping on *H. lupulus* reference, which is likely explained by the high value of $p$.

Furthermore, the authors of the *H. lupulus* genome assembly put forward the hypothesis of two Whole Genome Duplications (WGD) to explain the size of *H. lupulus* genome (>3Gb; Padgitt-Cobb *et al*., 2019). Although they filtered these duplications in the “deduplicated” annotation, a BUSCO analysis of the assembly revealed that 28% of genes are still duplicated (Padgitt-Cobb *et al*., 2019). The absence of sex-linked genes identification, the high value of *p*, and the high rate of duplicated genes in *H. lupulus* assembly convinced us to favour *C. sativa* reference transcriptome despite a lower mapping quality.

#### Expression results and ASE analysis

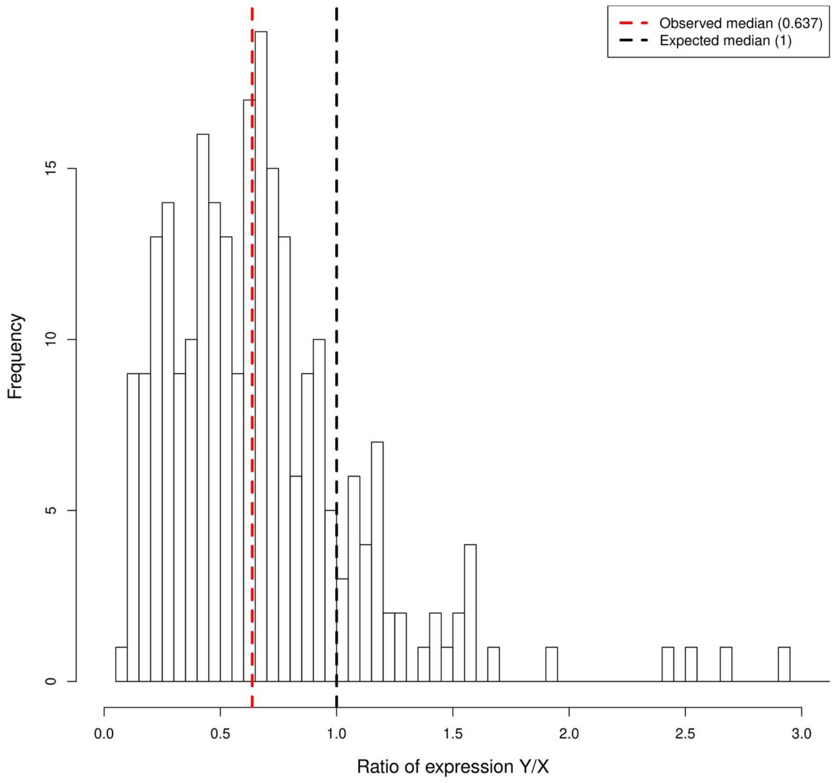


Figure S1. Histogram of the Y/X expression ratio. The dotted red bar represent the observed median (=0.637), and the dotted black bar the expected median of the ratio without Y degeneration (=1).

We quantified the ASE in the non-recombining region to determine the presence of X Chromosome Inactivation (XCI) in females. We computed the ratios of expression of one allele (A1) over the other one (A2) for each gene. We represented histograms of these ratios for genes in the non-recombining region and for the whole transcriptome. Figure S2 shows these histograms for the parents, Figure S3 for the daughters in the non-recombining region and Figure S4 for the daughters out of the non-recombining region.

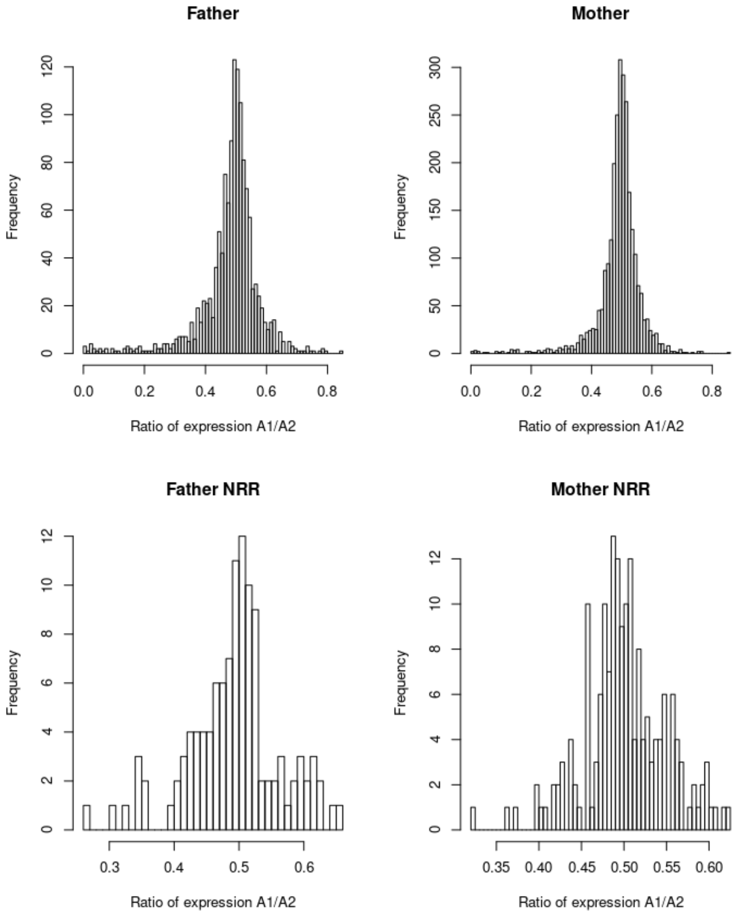


Figure S2. Histogram of ratio of A1 expression / A2 expression. Histograms are presented for the two parents. On the top, the ASE for the whole transcriptome, on the bottom, the ASE for the non recombining region (NRR) only.

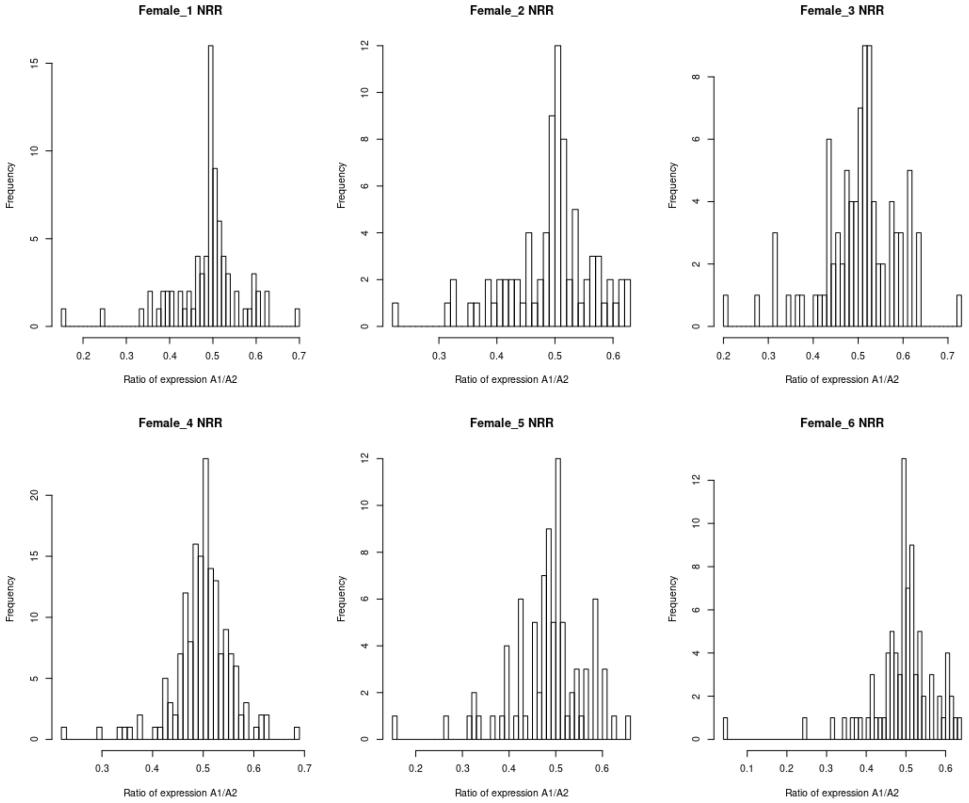


Figure S3. Histogram of ratio of A1 expression / A2 expression in daughters for genes in the non-recombining region (NRR).

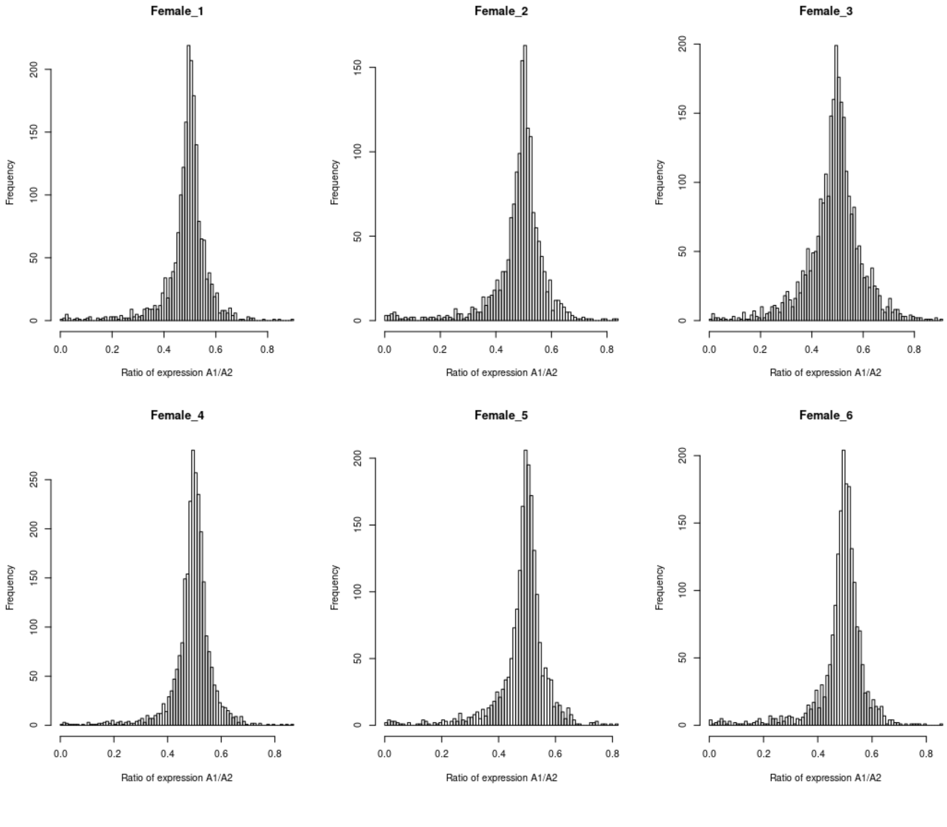


Figure S4. Histogram of ratio of A1 expression / A2 expression in daughters for genes out of the non-recombining region.

The three latter Figures don't support the hypothesis of an XCI since the mode of the allele expression ratio distribution is close to 0.5 in the non-recombining region. This ratio indicates that both alleles are expressed in same proportion, which is not expected with XCI.

#### Synonymous divergence with *M. notabilis* and *R. chinensis*

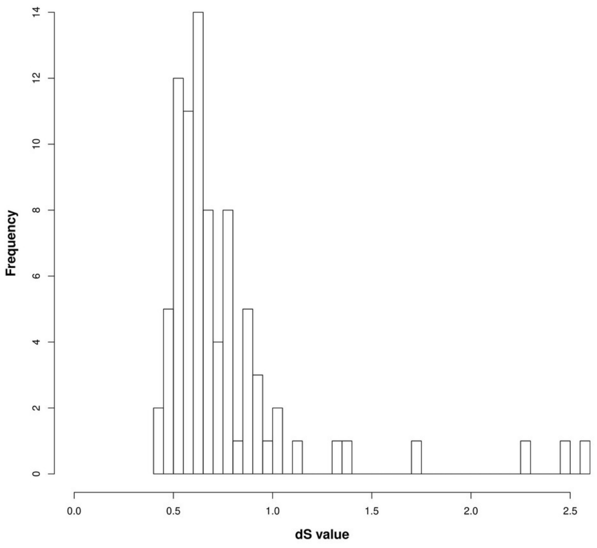


Figure S5. Histogram of synonymous divergence *(dS)* between *C. sativa* and *M. notabilis*. The lowest value is 0.437.

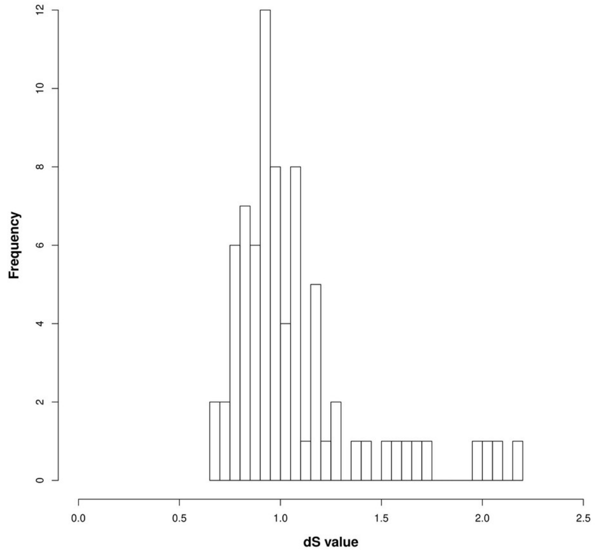


Figure S6. Histogram of synonymous divergence *(dS)* between *C. sativa* and *R. chinensis*. The lowest value is 0.673.

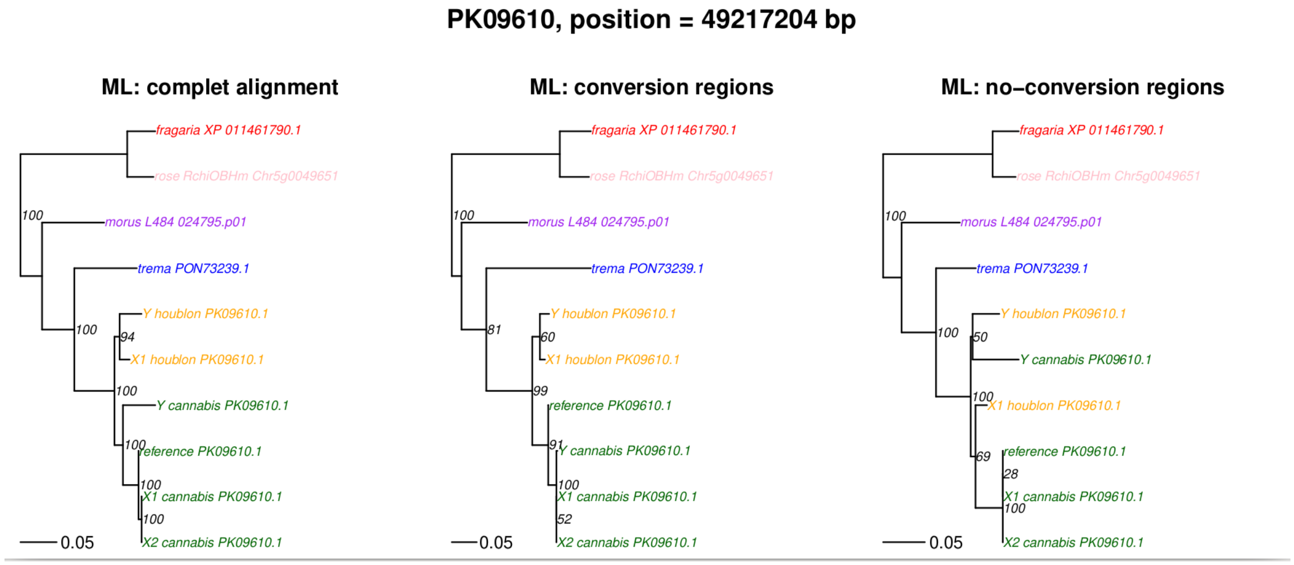


Figure S7. Phylogenetic results with maximum likelihood (ML) method for the gene PK11270 (on chromosome 1) for which geneconv identified gene conversion in a region representing around 50% of the gene length. On the left: the topology obtained for the whole sequence, in the middle: topology obtained for the region identified by geneconv as gene conversion, on the right: topology obtained for the region of the gene without gene conversion.

#### Reduction of Y expression and dosage compensation

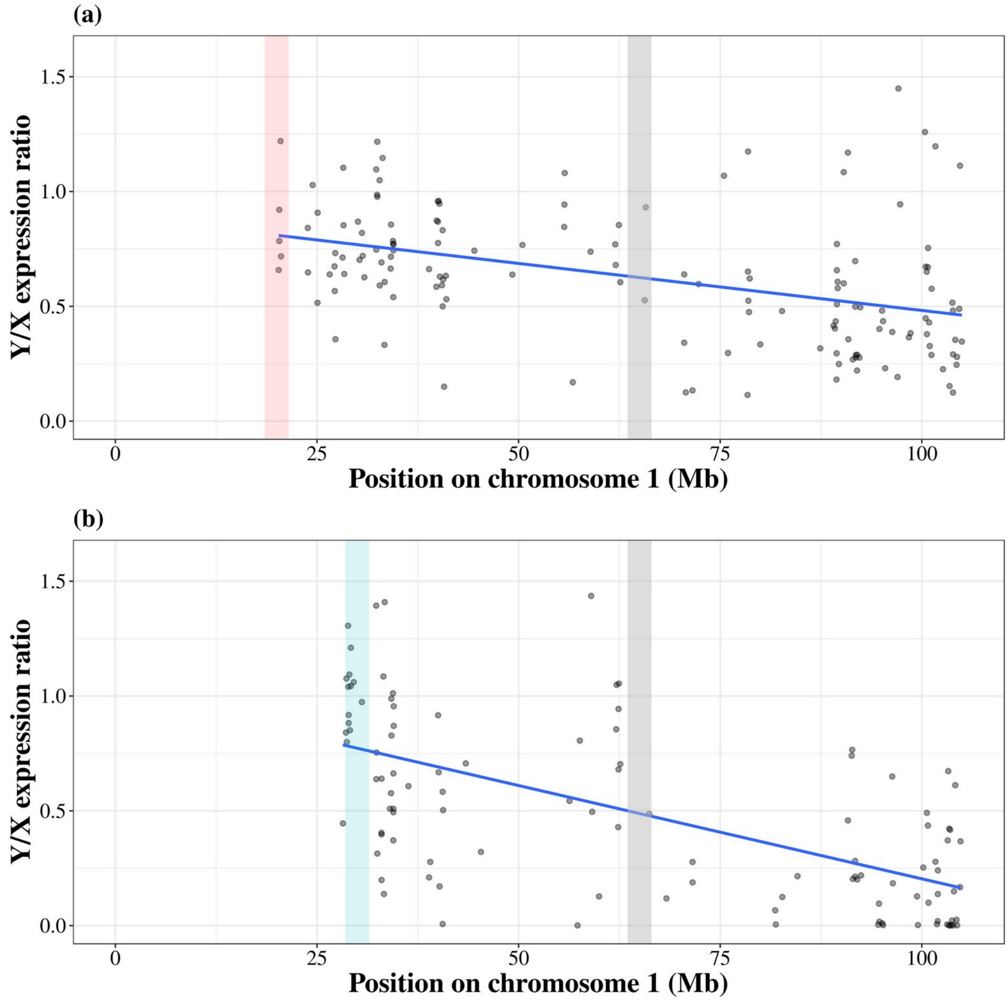

Figure S8. Y/X expression ratio along the X chromosome in *H. lupulus* **(a)**, and *C. sativa* **(b)** for genes without the detection of mapping bias with geneconv. The grey dots represent the Y/X expression ratio for each gene in the non-recombining region only. The blue line represents a linear regression (adjusted $R^2$=0.174, p-value<$10^{-5}$, and adjusted $R^2$=0.391, p-value<$10^{-5}$ for *H. lupulus* and *C. sativa*, respectively). The vertical red bar represents the putative Pseudo-Autosomal Boundary (PAB) in *H. lupulus*, the vertical blue bar represents the putative PAB in *C. sativa*, the vertical grey bar represents the putative boundary between the region that stopped recombining in the common ancestor and the region that stopped recombining independently in the two species.

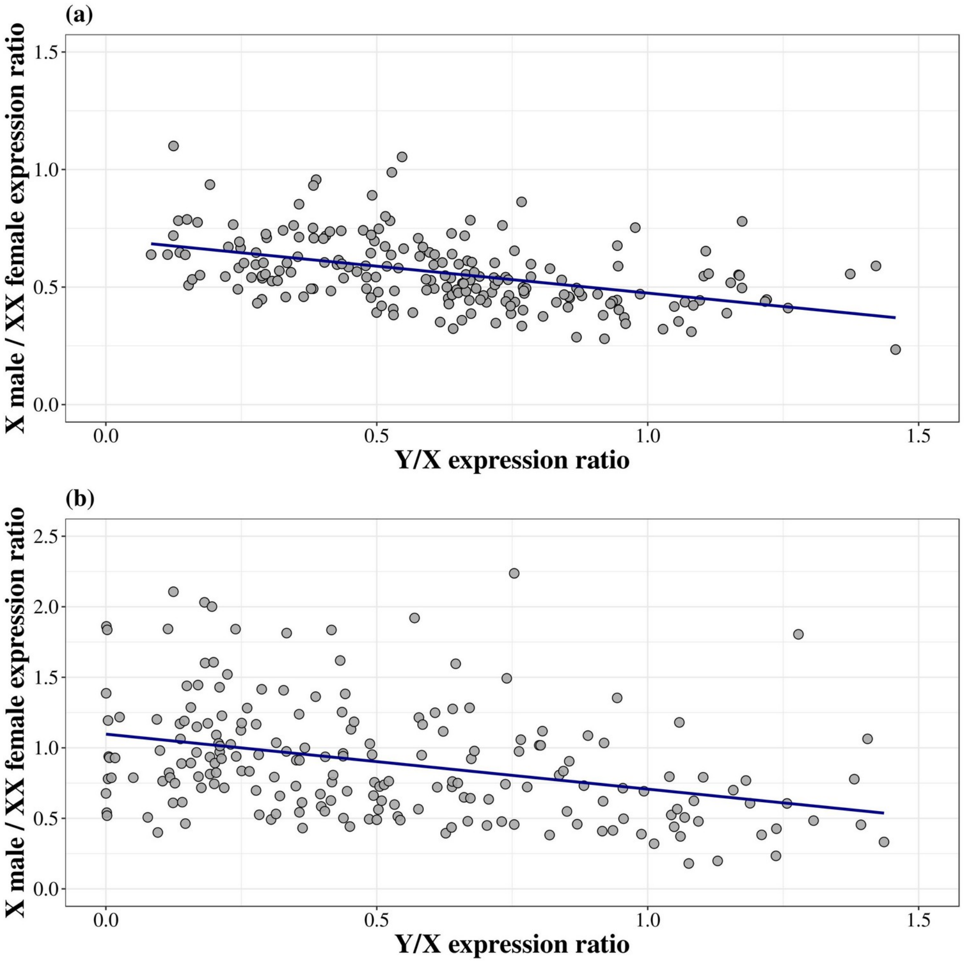


Figure S9. The male X expression over female XX expression versus Y/X expression ratio for *H. lupulus* (a) and *C. sativa* (b). Each black dot represents one gene. The blue line represents a linear regression (adjusted $R^2$=0.199, p-value<$10^{-5}$, and adjusted $R^2$=0.102, p-value<$10^{-5}$ for *H. lupulus* and *C. sativa*, respectively).

We repeated the expression level analyses using a dataset without the genes with possible mapping bias. The adjusted $R^2$ is always greater when removing genes for which we identified a mapping bias, as summarized in Table S3. This results confirmed that the reduction of the Y expression and the dosage compensation for genes with a Y expression strongly reduced, are not induced by the mapping bias of Y divergent sequences.

Table S3. Summary of adjusted $R^2$ for Y/X expression ratio and dosage compensation analyses for the two datasets. Dataset 1 = all genes; and Dataset 2 = genes without mapping bias identification only.

| | *H. lupulus* | | *C. sativa* | |
|---|---|---|---|---|
| | Dataset 1 | Dataset 2 | Dataset 1 | Dataset 2 |
| Dosage compensation analysis | 0.179 | 0.199 | 0.097 | 0.102 |
| Y/X expression ratio analysis | 0.134 | 0.174 | 0.278 | 0.391 |

#### Duplications impact on SEX-DETector inferences

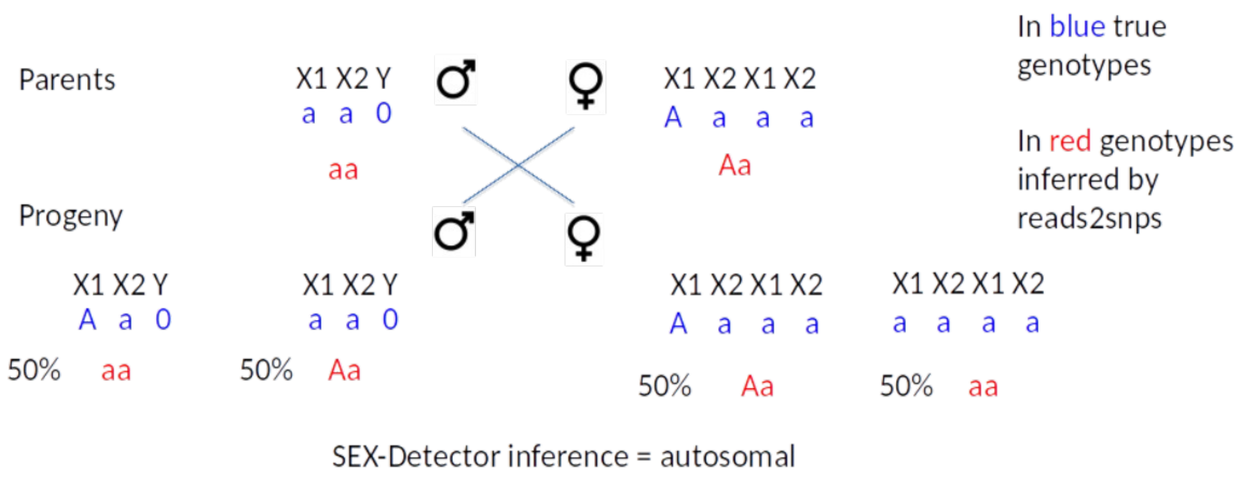


Figure S10: Hypothesis to explain the absence of X-hemizygous genes. If the *H. lupulus* X chromosome comprised two X1 + X2 chromosomes (from a possible WGD, see Padgitt-Cobb *et al*., 2019) whose reads map to the *C. sativa* X chromosome, then X-hemizygous genes will be impossible to detect. This is because ploidy differences between males and females is no longer detectable as shown in the example above. In this example, X-hemizygous genes will have an autosomal-like segregation. We should observe an excess of autosomal genes on the X-specific region XSR in *H. lupulus* (compared to *C. sativa* ). The X-hemizygous genes detected in *C. sativa* should have a XY or autosomal segregation type in *H. lupulus.*
